# Design Principles of the ESCRT-III Vps24-Vps2 Module

**DOI:** 10.1101/2020.12.28.424573

**Authors:** Sudeep Banjade, Yousuf H. Shah, Shaogeng Tang, Scott D. Emr

## Abstract

ESCRT-III polymerization is required for all ESCRT-dependent events in the cell. However, the relative contributions of the eight ESCRT-III subunits differ between each process. The minimal features of ESCRT-III proteins necessary for function, and the role for the multiple ESCRT-III subunits remain unclear. To identify essential features of ESCRT-III subunits, we previously studied the polymerization mechanisms of two ESCRT-III subunits Snf7 and Vps24, identifying the association of the helix-4 region of Snf7 with the helix-1 region of Vps24 (Banjade et al., 2019). Here, we find that mutations in the helix-1 region of another ESCRT-III subunit Vps2 can functionally replace Vps24 in *S. cerevisiae*. Engineering and genetic selections revealed the required features of both subunits. Our data allow us to propose three minimal features required for ESCRT-III function – spiral formation, lateral association of the spirals through heteropolymerization, and binding to the AAA+ ATPase Vps4 for dynamic remodeling.

## Introduction

ESCRTs (endosomal sorting complexes required for transport) control a growing list of membrane remodeling events in cells (*1*). Among the different subcomplexes of ESCRTs, ESCRT-III is required in all processes, while the requirement of the other upstream complexes 0, I and II is variable (*2*). Our understanding of the mechanisms of ESCRT-III assembly has increased substantially in the last decade (*3–17*). However, important questions remain. The specific role of each of the eight ESCRT-III subunits remain unclear. In eukaryotes, eight ESCRT-III proteins exist: Did2 (CHMP1), Vps2 (CHMP2), Vps24 (CHMP3), Snf7 (CHMP4), Vps60 (CHMP5), Vps20 (CHMP6), Chm7 (CHMP7) and Ist1 (IST1) (*18, 19*). Of these, Vps20, Snf7, Vps24 and Vps2 have been studied in greater detail, because they are individually essential for MVB biogenesis in yeast, the earliest ascribed role for ESCRTs. These four proteins are the minimal subunits necessary to create intraluminal vesicles in the MVB pathway, as suggested by *in vivo* and *in vitro* analyses (*9, 11, 17, 20*). A recent study has further included Did2 and Ist1, providing additional insights into the sequential recruitment mechanism of ESCRT-III components (*17*). Toward a comprehensive understanding of each ESCRT-III subunit, studies in MVB biogenesis have provided important clues regarding their individual roles in membrane remodeling.

Vps24 and Vps2 are essential for MVB biogenesis (*3, 21, 22*). Vps24 and Vps2 are also recruited cooperatively to membranes, requiring each one for the other’s efficient recruitment (*9, 10, 21, 23*). Because of this reason, these two proteins have been analyzed together in previous ESCRT related work. Interestingly, in previous work, it was found that during HIV budding, while CHMP2 (the human ortholog of Vps2) is essential, CHMP3 (the human Vps24 ortholog) is not essential for HIV egress from cells (*24*). We hypothesized that there may be features of Vps24 and Vps2 that renders CHMP3 non-essential in some ESCRT-dependent processes. What are these essential features in these two proteins that make them indispensable for membrane remodeling? We set out to define those features in this work.

One of the most important features of Vps2 that has previously been described is the recruitment of the AAA+ ATPase Vps4, which modifies ESCRT-III polymers(*25, 26*). Vps24 and Vps2 bind to Snf7 and remodel Snf7 polymerization, changing the flat Snf7 spiral into 3D helices (*3, 15, 17, 23*). We hypothesized that a single protein with features of Vps24 and Vps2 that can bind to Snf7 but also recruit Vps4 may be sufficient for function. Here, through engineering approaches we define a single protein that possesses such functions of both Vps24 and Vps2. These data allow us to define the minimal essential properties of an ESCRT-III heteropolymer that are required for intraluminal vesicle formation. These include - a) spiral formation through a Snf7-like molecule, b) lateral association through a Vps24/Vps2 like molecule and c) the ability to recruit the AAA+ ATPase Vps4.

## Results

### Overexpressing Vps2 can replace the function of Vps24 in MVB sorting

In our previous work (*23*), we observed that overexpressing *VPS24* suppresses the defect of a *snf7* allele (*snf7-D131K*) that encodes a Snf7 mutant with a lower affinity to Vps24. The overexpression, however, does not rescue other *snf7* alleles that encode defective Snf7 homo-polymers. We also observed that overexpression of Vps2 rescues the defect of *snf7-D131K*. These data are consistent with the observations that Vps24 and Vps2 bind synergistically to Snf7 (*9, 10, 21*). Following these observations, we sought to test whether expressing a high level of Vps2 also rescued the lack of Vps24 in cells, with the hypothesis that Vps2 may possess a lower affinity binding surface for Snf7, which could be overcome with an increased availability of Vps2 in the cytoplasm.

To test these hypotheses, we first utilized MVB cargo sorting assays (for cargoes Mup1 and Can1) in *S. cerevisiae (23, 27*). In a *vps24*Δ strain overexpressing *VPS2*, we observed that Mup1 −pHluorin is sorted at about 40% compared to that of the wild type, and that the canavanine sensitivity of *vps24*Δ is partially rescued (Figure 1A). We also noted that *VPS2* overexpression rescues the temperature sensitivity of *vps24*Δ (Figure 1 – Fig. Supp 1A), suggesting that the increased concentration of Vps2 could replace the cellular function of Vps24 beyond MVB cargo sorting.

**Figure 1.**
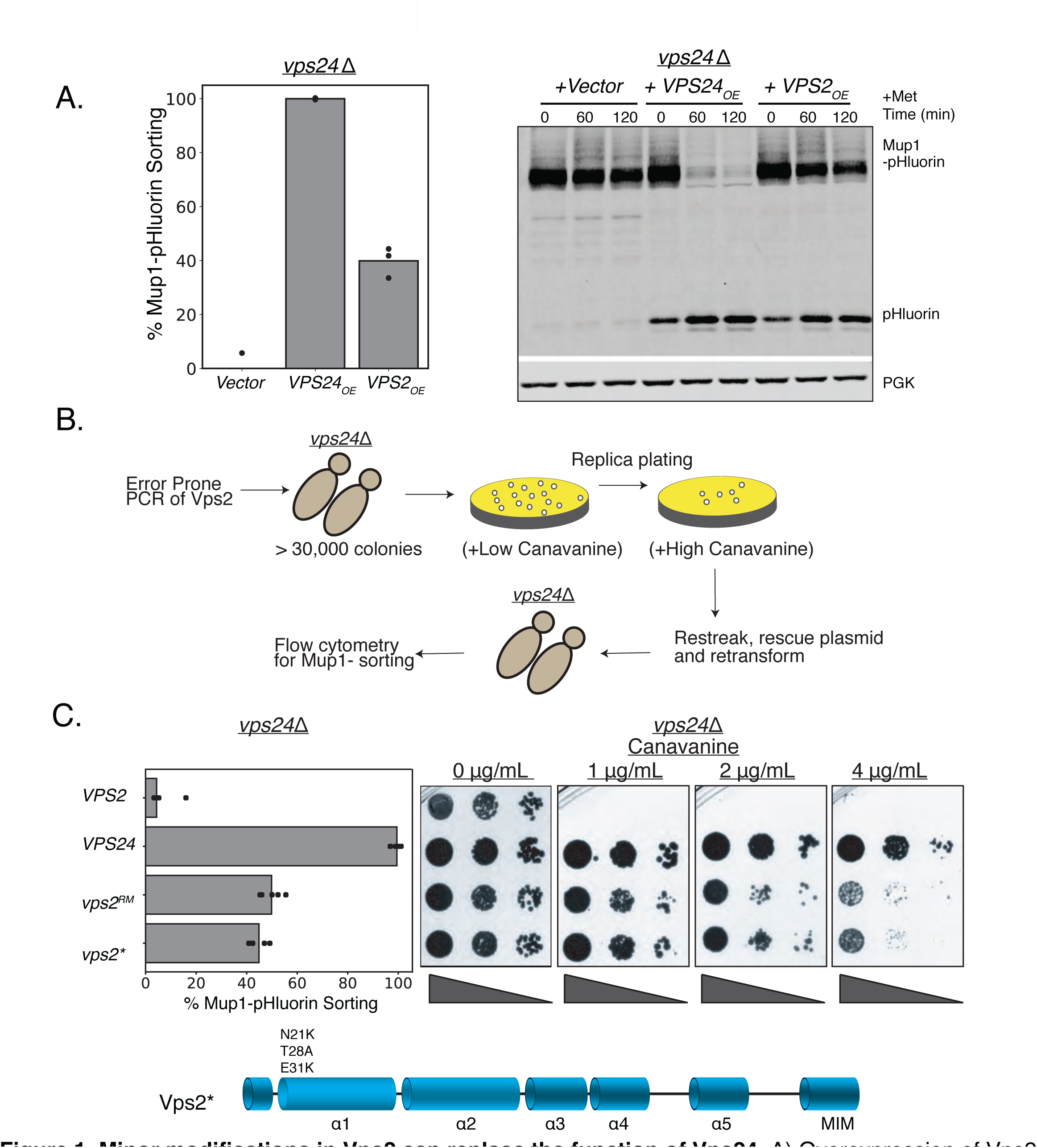
Minor modifications in Vps2 can replace the function of Vps24. A) Overexpression of Vps2 can rescue the defect of *vps24*Δ for Mup1 sorting. Image on the left represents Mup-pHluorin sorting through a flow-cytometry assay and the image on the right represents an immunoblot for pHluorin upon methionine addition. Overexpression (OE) was achieved through a CMV-promoter and Tet-operator containing plasmid. B)Flow-chart of the random mutagenesis approach. C) Top figure shows the flow cytometry and canavanine sensitivity assays with the mutant Vps2 that can rescue the sorting defects of *vps24*Δ. Bottom figure shows the domains of Vps2 highlighting the mutations found in the selection.

With a tet-off regulatable operator, we next used doxycycline to titrate the expression level of Vps2. As a result, we determined that about 8-fold overexpression of Vps2 is necessary for restoring Mup1 sorting (Figure 1 – Fig. Supp 1B-D). These data suggested that Vps2 contains features that can replace the function of Vps24 when present in higher concentrations in cells.

### Random mutagenesis and selection of *vps2* mutants that are capable of replacing both *VPS2* and *VPS24*

To identify the features in Vps2 that could replace the function of Vps24, we utilized an unbiased random-mutagenesis selection approach, as we have done previously (*23, 28*). Since *vps24*Δ is sensitive to the drug canavanine, we selected for *vps2* mutants that conferred canavanine resistance to *vps24*Δ cells. We performed error-prone PCR and assembled a *vps2* mutant library in a *vps24*Δ strain. We next selected *vps2* alleles using canavanine at a concentration that the wild-type *VPS2* does not grow (Figure 1B). From this selection, one of the alleles (hereafter referred to as Vps2^RM^) strongly rescues the canavanine sensitivity of *vps24*Δ (Figure 1C), and sorts Mup1-pHluorin to 45% of that of the wild-type (Figure 1C). Vps2^RM^ contains mutations in its promoter region, three missense mutations in helix-1 (N21K T28A E31K), and two missense mutations in helix-4 (S136N M146I).

The N-terminal mutations E21K T28A N31K (with the promoter mutations, hereafter called Vps2*) are necessary and sufficient for the suppression effect. Interestingly, these mutants also lie on the same surface of the alpha-1 helix, as a helical-wheel representation suggests (Fig. 1 – Figure Supp. 3B). We found that while the individual mutations did not rescue *vps24*Δ (Figure 1 – Fig. Supp. 2A), they collectively suppressed both the defect in canavanine sensitivity, and Mup1 (sorting upto ~40%). Because the mutant we isolated also had promoter mutations, the expression level of Vps2* is increased about 3-fold (Figure 1 – Fig. Supp 2C). The suppression by Vps2* is not due simply to the overexpression effect however, since about 8-fold overexpression of Vps2 is required for only ~20% sorting of Mup1 to occur in a *vps24*Δ strain (Figure 1 – Fig. Supp 1A-C).

To test whether Vps2* possesses features of both Vps24 and Vps2, we performed cargo sorting assays in *vps24*Δ*vps2*Δ. We observed that *vps2\** also suppresses *vps24*Δ*vps2*Δ (Figure 2A-B). Therefore, a synergistic effect of the mutations in the N-terminal basic region of Vps2 helix-1 and its 3-fold overexpression provide the necessary functional features of Vps24 and Vps2.

**Figure 2.**
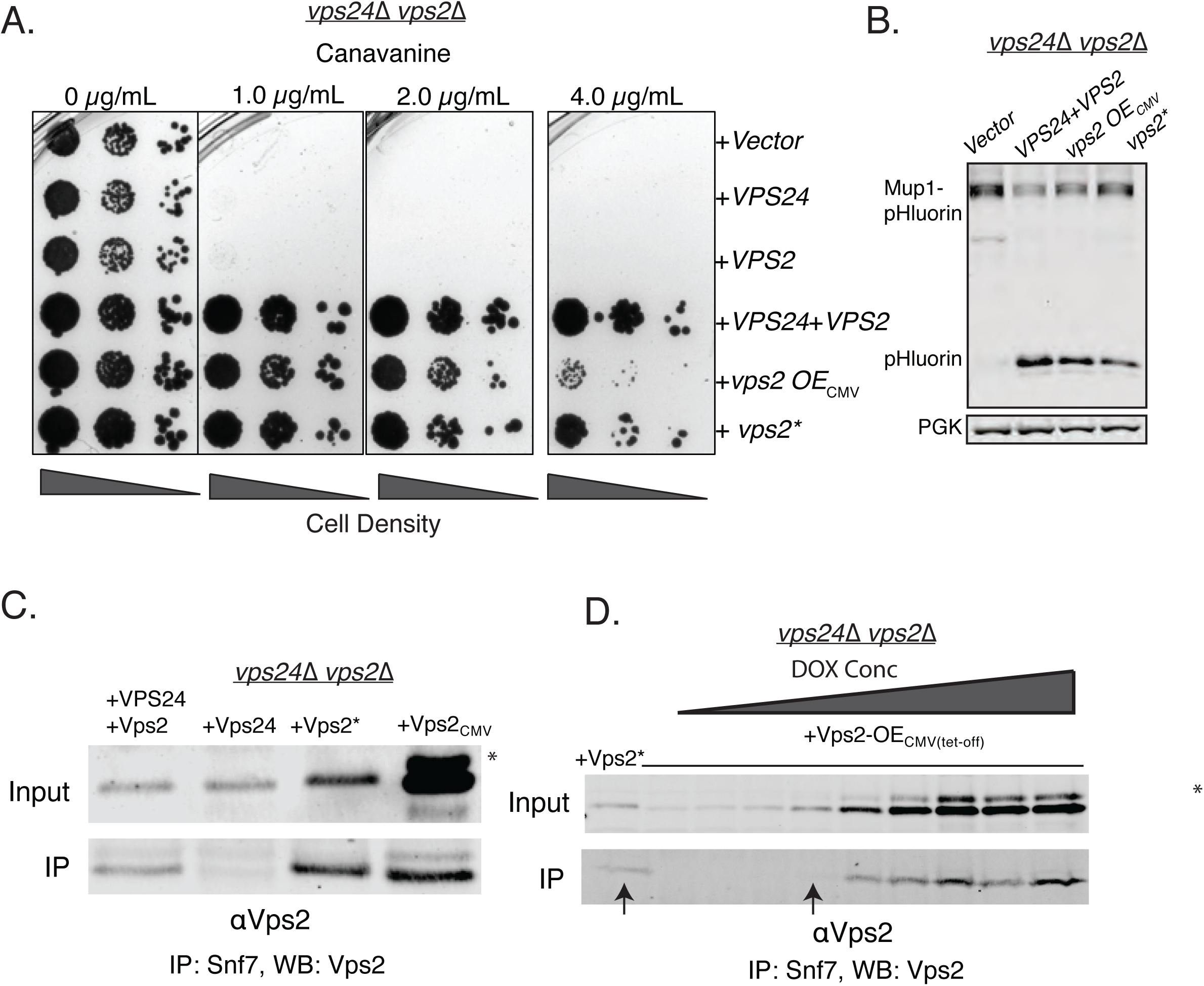
Properties of both Vps2 and Vps24 in a single Vps2 contruct. A) Canavanine sensitivity data in *vps24*Δ *vps2*Δ with an overexpression of Vps2 (CMV-Tet system) or with Vps2*. B) Immunoblot for Mup1-pHluorin sorting upon overexpression of Vps2 (CMV) or with Vps2*. C) Co-immunoprecipitation of Snf7 with Vps2 (CMV) and Vps2*. D) Coimmunoprecipation experiments of Snf7 with Vps2 at various expression levels of Vps2 after titration of the tet-off operator with doxycycline. Arrows point to the relative binding to Snf at similar expression levels of Vps2 and Vps2*. In the gels, * refers to an unknown modified form of Vps2.

One of the early-identified functions of Vps24 in yeast was as an adaptor for Vps2 to be recruited to Snf7 polymers (*20*). In the absence of Vps24, Vps2 does not bind to Snf7 in coIP experiments (Figure 2C and as observed in (*21*),). This effect is rescued by the CMV promoter-mediated overexpression of Vps2 or with Vps2*.

Vps2* binds to Snf7 at a lower expression level than the WT Vps2, providing further evidence that the N-terminal helix-1 mutations increase the affinity of Vps2 for Snf7, bypassing the need for Vps24 (Figure 2D). Although the overall feature of helix-1 region of Vps24 and Vps2 are similar (both basic helices), they vary in sequence composition (Figure 1 – Fig. Supp. 3A). Since the mutations occur in polar residues (Figure 1 – Fig. Supp. 3A-B), it is possible that these charge inversions increase the affinity of the basic patch of Vps2* to Snf7’s acidic helix-4, consistent with our observations of Vps24 binding to Snf7 through an electrostatic interface (*23*). Simple overexpression of Vps2 probably rescues the defect of the lack of Vps24 since the overall concentration of a lower-affinity Snf7-binding Vps2 molecule is increased in the cell.

### Binding to the AAA+ ATPase Vps4 is a critical feature of the Vps24-Vps2 module

In contrast to Vps2 overexpression rescuing the defects of *vps24*Δ, the reverse doesn’t occur – Vps24 overexpression by ~16 fold did not rescue the defect of a *vps2*Δ (Fig. 3). One of the critical features of Vps2 is the presence of the C-terminal MIM motif that has a higher affinity to the AAA+ ATPase Vps4 than other ESCRT-III proteins (*25*). While other ESCRT-III proteins also possess the MIM motifs, the Vps2 MIM has the strongest affinity for Vps4 in solution (~20 μM, (*25*)). In addition, helix-5 of Vps2 has been identified as a second binding site of Vps4 with an affinity of ~3 μM (*29–31*). We therefore hypothesized that a Vps24 variant with Vps4 binding sites could possesses the properties of both Vps24 and Vps2. To test this, we replaced the C-terminus of Vps24 with that of Vps2 and assayed the functions of the chimeric protein.

**Figure 3.**
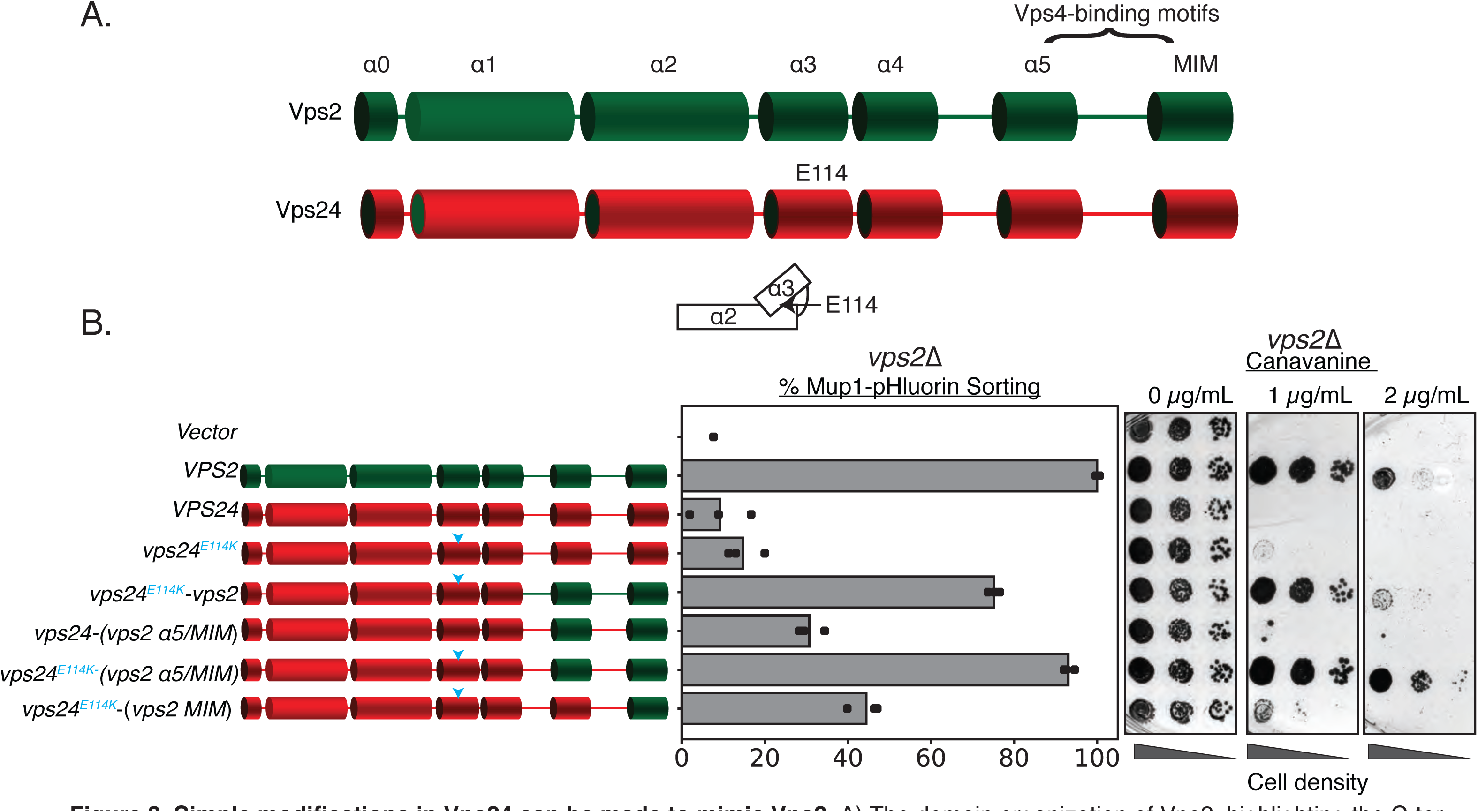
Simple modifications in Vps24 can be made to mimic Vps2. A) The domain organization of Vps2, highlighting the C-terminal region important for Vps4 binding. B) Left-panel denotes the chimeras made to replace regions of Vps2 onto Vps24. Cyan arrows in the helices are positions of the E114K mutation. Right panel represents Mup1-pHluorin sorting and canavanine-sensitivity assays. In this assay, the constructs were over-expressed under a CMV-promoter-Tetoff operator system, overexpressing Vps24 ~16 fold.

We designed various chimeric constructs of Vps24/Vps2 under the control of their endogenous promoters to first demarcate the regions that maintain function in a *vps24*Δ strain (Figure 3 – Fig. Supp 1A). Consistent with the sequence analysis, replacing the MIM and helix5 of Vps24 with the homologous region of Vps2 kept the protein functional, but truncations beyond residue ~152 resulted in functional defects. In summary, the C-terminus (residues ~152 and beyond) of Vps24 can be replaced with that of Vps2 and still retain function.

We next tested these chimeras in *vps2*Δ. In contrast to *vps24*Δ cells, we observed that they do not functionally restore the loss of Vps2 (Figure 3 – Fig. Supp 1B). To investigate whether they are dependent on protein expression levels, we then overexpressed these constructs with a CMV promoter that contain the N-terminus of Vps24 and helix-5 and MIM of Vps2. We found that they modestly suppress the defect of *vps2*Δ to about 30% of wild-type (Figure 3). Therefore, directly recruiting Vps4 on to Vps24 partially bypasses the requirement of Vps2. We speculated that there are additional features in Vps24 that make it distinct from Vps2.

ESCRT-III subunits are soluble monomers in the cytoplasm but undergo structural rearrangements when assembled on membranes. We next investigated whether activating mutations in Vps24 that induce conformational changes renders it similar to Vps2. In our previous work (*28*), we designed a Snf7 mutant (hereafter referred to as Snf7***) that rescues the defects of *vps20*Δ. Snf7*** includes a myristolyation site that recruits Snf7 to endosomes in the absence of upstream factors (ESCRT-I, ESCRT-II and Vps20) (*28*), as well as missense mutations (R52E Q90L N100I) that trigger Snf7 to adopt an elongated, open, and membrane-bound conformation (*2, 22*).

Inspired by our early studies, we looked for single amino acid substitutions in Vps24 that weaken the interactions between helix-3 and helix-2 that would allow an extension of helices-2 and 3, and therefore “activate” Vps24. As a result, we found that *vps24^EII4K^* when overexpressed robustly rescues the defect of *vps2*Δ (Figure 3). Remarkably, this rescue becomes more pronounced when *vps24^EII4K^* carries the Vps4-binding motifs of Vps2 (Figure 3). Taken together, our data suggest that a Vps24 variant capable of auto-activation and Vps4 recruitment can function as Vps2.

### Conformationally distinct Vps24 and Vps2 species replace one another

In the “closed” conformation of ESCRT-III subunits, the region from helix 3 and beyond bind back to alpha helices 1-2 (*8, 19, 32*). During activation, the extension of helices 2-3 into an elongated helix triggers “opening” of the protein, which enables polymerization due to the availability of an extended surface for self-assembly. Therefore, mutations that trigger conformational changes to an “open” state are able to readily form polymers *in vivo* and *in vitro*. Consistent with these ideas, Vps24^E114K^ in cell lysates forms higher-molecular weight species in glycerol-gradient experiments (Figure 4A). In *in-vitro* assays, while the wild-type Vps24 doesn’t form polymers by itself or with Vps2, Vps24^E114K^ readily associates into linear filaments with Vps2 at concentrations of 5 μM each (Figure 4B, Figure 4 – Fig. Supp. 1A). In comparison, previous experiments with Vps24^WT^ required 70 fold higher concentrations to observe similar linear polymers (*33*). With Snf7 and Vps2, both wild-type and the mutant Vps24 are able to form 3D spirals, which we previously described as the copolymeric structure of Snf7, Vps24 and Vps2 (Figure 4 – Fig. Supp. 1B). The increased ability to form polymers is consistent with the interpretation that E114K shifts the equilibrium of Vps24 to the open state, and that this open state mimics Vps2.

**Figure 4.**
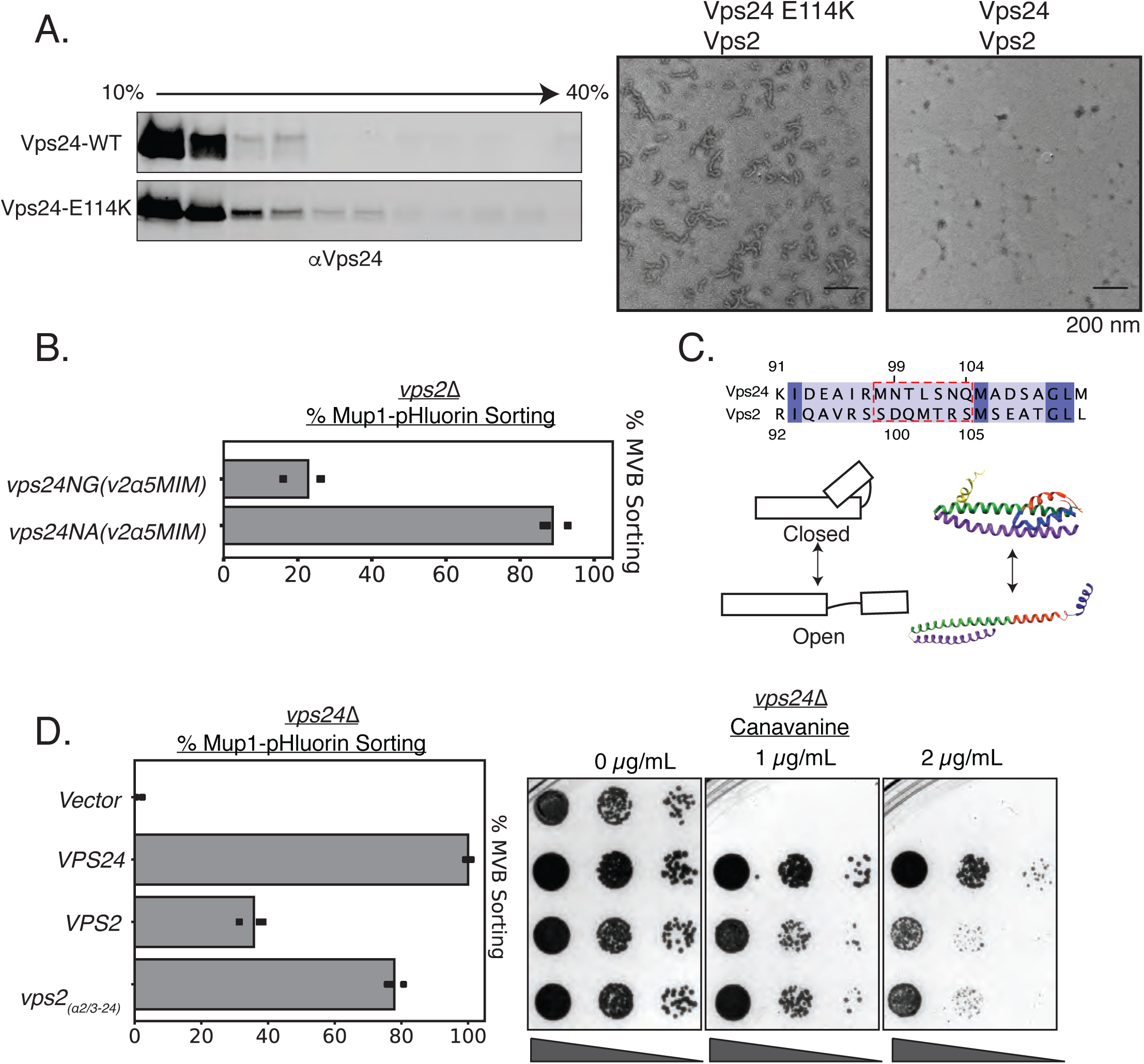
Vps24 and Vps2 may exhibit different conformations. A) Left: glycerol-gradient experiments with Vps24 and Vps24E114K suggests that the mutant can form higher molecular-weight species. Right: negative stain electron microscopy of Vps24 E114K or WT Vps24 at 1 *μ*M each of the proteins in the presence of Vps2. B) Mup1-pHluorin assays with Vps24 mutations in the asparagines (N99 and N103) α2/α3 hinge region to Ala or Gly residues in constructs that have the Vps4-binding sites H5(helix-5) and MIM from Vps2 (V2). These constructs are expressed with the CMV-promoter, Tet-off system.C) Top: sequences of the α2/α3 hinge region of Vps24 and Vps2. Bottom left: Model showing the two conformations of ESCRT-III proteins. Structural model on the right is that of CHMP3(closed) (Bajorek, 2009) and of Snf7(open) (Tang, 2015). D) Mup1-pHluorin sorting and canavanine-sensitivity assays with overexpression of Vps2 (CMV-Tet) and with a mutant replacing the α2/α3 hinge region of Vps2 with that of Vps24 (also CMV-Tet system).

From sequence alignment analysis (Fig. 4C), we noticed low conservation between Vps24 and Vps2 in the hinge between α2 and α3. Vps24 contains two helix-breaking asparagine residues in between these alpha helices (N99 and N103), while Vps2 lacks these helix-breaking residues (Figure 4C). We found that mutating the Asn to helix-stabilizing Ala (in addition to the Vps4 binding motifs) in Vps24 rescues the defect of *vps2*Δ, while the helix-breaking glycine does not rescue the defect (Figure 4B).

Therefore, it appears that the hinge region, that contains residues which affect the conformational flexibility of ESCRT-III proteins, in addition to the Vps4 binding sites, accounts for the majority of the difference between Vps24 and Vps2. Consistent with this, when we overexpress a variant that replaces the hinge region of Vps2 with that of Vps24, it suppresses *vps24*Δ to ~80% as tested by Mup1 sorting, compared to ~40% for that of the wild-type Vps2 (Figure 4D).

These data may suggest that Vps24 may not possess the ability to fully extend its helices-2 and 3 in the ESCRT-III copolymer. A recently published CryoEM structure showed that the Vps24 homopolymer consists of Vps24 protomers in a “semi-open” conformation (*34*) (see Fig. 4 – Fig. Supp. 2 for direct comparison), in contrast to the fully extended and open Snf7 (*8*) and Did2 (CHMP1) (*16*) polymers. It is possible that a mixture of different conformations allows for efficient Vps24-Vps2 assembly, which has a higher affinity to the Snf7 polymer.

Collectively, these data suggest that while Vps24 and Vps2 are similar proteins. Laterally interacting with Snf7, inducing the formation of an ESCRT-III super-helix, and recruiting Vps4 are three features for the Vps24/Vps2 module.

### “Accessory” ESCRT-III genes promote intralumenal vesicle formation

In *Saccharomyces cerevisiae*, there are eight ESCRT-III genes. One of the defining features of these ESCRT-III proteins is the N-terminal alpha-helical bundle, which is sometimes referred to as the ESCRT-III domain. The other defining feature is the C-terminal flexible region that contains at least one MIM, that binds to the MIT domain of Vps4. The N-terminus of the ESCRT-III domain are similar in sequence and structure. However, the specific functions of all these ESCRT-III proteins remains unclear.

To quantitatively assess the relative contributions among the ESCRT proteins, we assayed for Mup1-pHluorin sorting in each gene deletion (Figure 5– Fig. Supp 1). We also performed canavanine sensitivity assays (Fig. 5 – Fig. Supp 1 B) with the same mutants. We observed that *snf7*Δ, *vps20Δ, vps24*Δ, and *vps2*Δ show severe sorting defects, *did2*Δ and *vps60*Δ show partial sorting defects, and *ist1*Δ and *chm7*Δ show no defect. These data are consistent with previous findings with a different cargo (CPS) (*18*), and from in-vitro assays (*11, 17*), which suggest that Snf7, Vps20, Vps24 and Vps2 are the minimal contributors in MVB formation.

**Figure 5.**
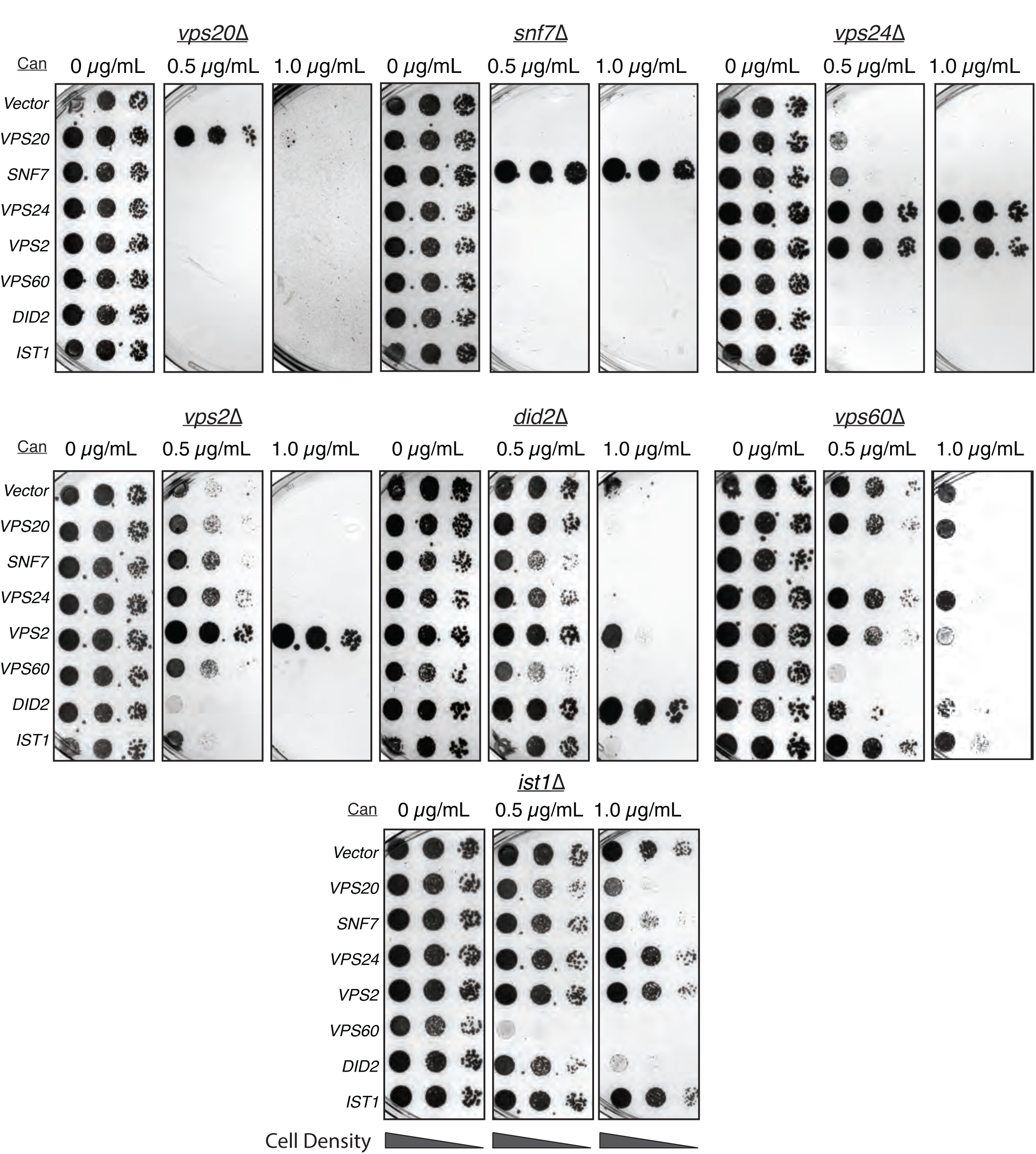
Overexpressing ESCRT-III proteins in the background of other ESCRT-III mutants show selective rescue phenotypes. In the annotated mutants, ESCRT-III proteins were expressed with a CMV-promoter/Tet-operator system, and plated in canavanine-containing plates. Vps2 overexpression can rescue the defect of *vps24*Δ. Vps2 overexpression in a *did2*Δ partially rescues canavanine sensitivity. Vps60 overexpression appears to be dominant negative.

These differences in function during MVB formation occur despite similarity in structure and sequence between these ESCRT-III proteins. Inspired by the observation that *VPS2* overexpression suppresses *vps24*Δ, we investigated whether overexpressing other ESCRT-III genes could suppress the deletions of a different ESCRT-III gene. We used the CMV promoter to overexpress each of the ESCRT-III genes: Snf7 is overexpressed by ~5 fold, and Vps24 and Vps2 by ~16 fold. Most of the overexpression constructs didn’t rescue the defect of the other ESCRT-III deletions, except in two cases. As described above, Vps2 overexpression rescued the defect of *vps24*Δ, and also partially rescued the defect of a *did2*Δ.

Evolutionary analyses have grouped ESCRT-III into two groups: Snf7/Vps20/Vps60, and Vps24/Vps2/Did2, (Fig. 6A) (*35, 36*). Vps20 nucleates formation of Snf7 spirals, and Vps24/Vps2 induce bundling and helix formation of spirals (*23*). In *in-vitro* assays with lipid monolayers, we found that Did2 forms tube-like helices (Fig. 6B, and as previously shown in (*7, 16*) for mammalian Did2 named CHMP3). However, Vps60 lacks the ability to form long helices/tubes, and preferentially forms spiral-like structures, reminiscent of Snf7 (Fig. 6B). Consistent with Vps60 mimicking Snf7 structurally, the N-terminal region of Snf7 fused to the C-terminal region of Vps60 rescues the defects of *vps60*Δ (Fig. 6C). Vps60-GFP is localized to endosomal and vacuolar membranes with a hint of plasma membrane signal (Fig. 6 – Figure Supp. 1). This localization is primarily cytosolic in *vps20Δ, snf7*Δ or *vps2*Δ, and unchanged in *did2*Δ (Fig. 6 – Figure Supp. 1).

**Figure 6.**
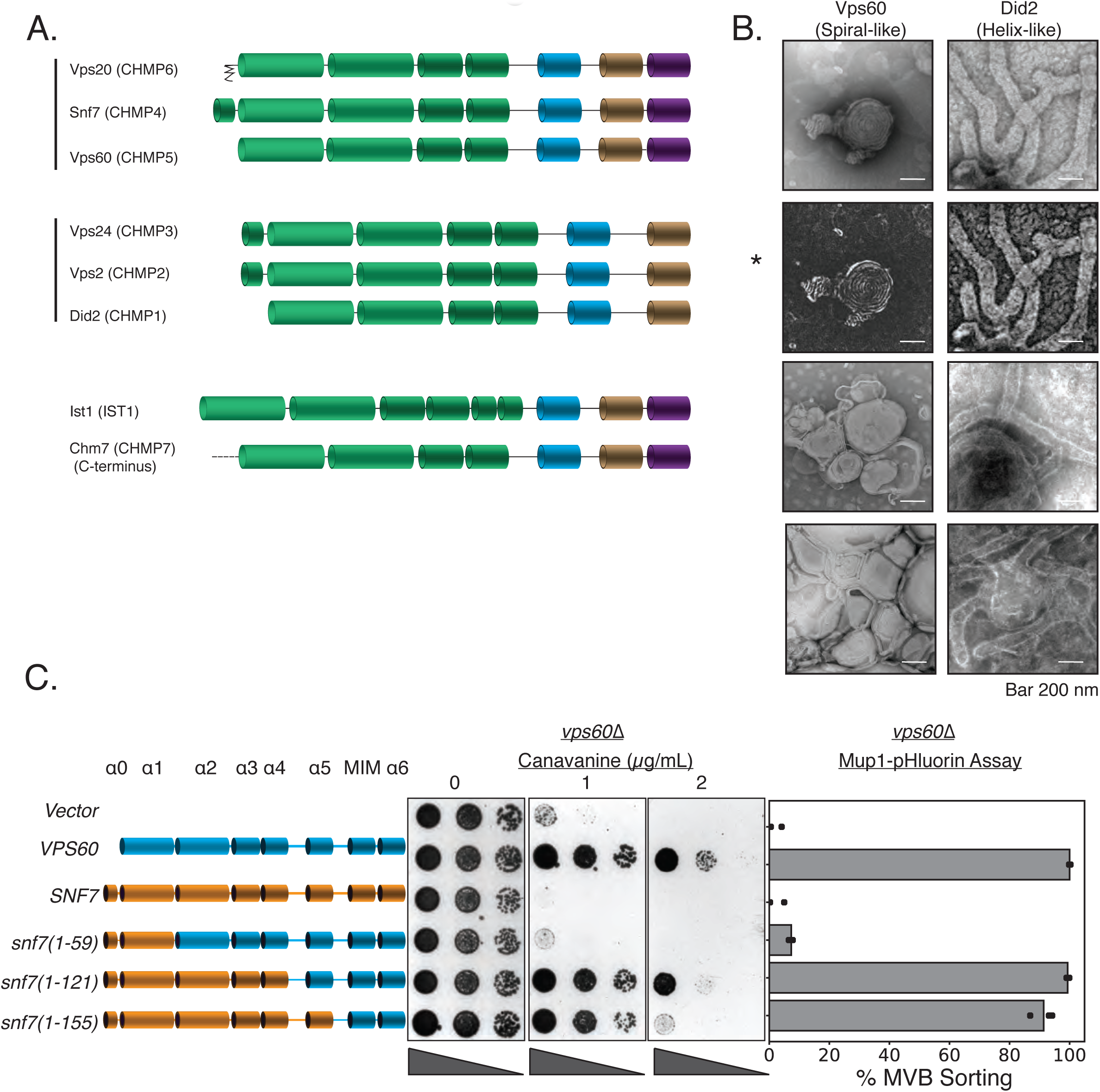
Vps60 possesses features of Snf7. A) Domain subunits of the eight ESCRT-III proteins in yeast. The mammalian names are in parentheses. B) Electron microscopy images of 1 *μ*M Vps60 or 1 *μ*M Did2 on lipid monolayers, incubated for 1 hour. Bar is 200 nm each.Top two images are different constrast-adjusted depictions of the same image. C) Domain swaps from Snf7 onto Vps60 can rescue the defects of canavanine senstivity and Mup1-pHluorin sorting in a *vps60*Δ strain.

In summary, among Vps20/Snf7/Vps20, Snf7 serves as the main scaffold which can be engineered to substitute for Vps20 (*28*) or Vps60 (Fig. 6A). Among Vps24/Vps2/Did2, modifications within Vps24/Vps2 can functionally replace each other; Did2 resembles Vps24/Vps2 as it readily forms three-dimensional helices, and *did2*Δ can be partially suppressed by Vps2 overexpression. Our data suggest that although ESCRT-III subunits have evolved for divergent roles in ordered assembly, rational modifications in ESCRT-III subunits can allow one to consolidate the functions of two ESCRT-III proteins into one ESCRT-III protein.

## Discussion

In this work we have utilized rational design and unbiased mutagenesis to understand the design principles of ESCRT-III subunits. To simplify this larger question, we focused primarily on the ESCRT-III subunits Vps24 and Vps2. First, we find that overexpression of Vps2, by ~8-fold and above, can rescue *vps24*Δ in yeast. Second, point mutations in the helix-1 region of Vps2 can also rescue *vps24*Δ. These Vps2 mutants also bind to Snf7 *in-vivo* even in the absence of Vps24. Third, overexpression and inclusion of higher-affinity AAA+ ATPase Vps4 binding regions on Vps24 can rescue the absence of Vps2. Fourth, mutations that induce conformational changes in Vps24 and Vps2 also rescue each other’s function. These data indicate a strong similarity in between these two ESCRT-III subunits.

Despite the observed similarity between Vps24 and Vps2, our data also suggest some differences: they may exist in different conformations (Fig. 7A), and that Vps2 consists of a higher-affinity Vps4 binding site (Fig. 7B). In addition, Vps24/Vps2 induce lateral association and bundling (*10, 17, 23*), along with helicity of the spirals, which could be an important parameter for ESCRT-III function for MVB biogenesis.

**Figure 7.**
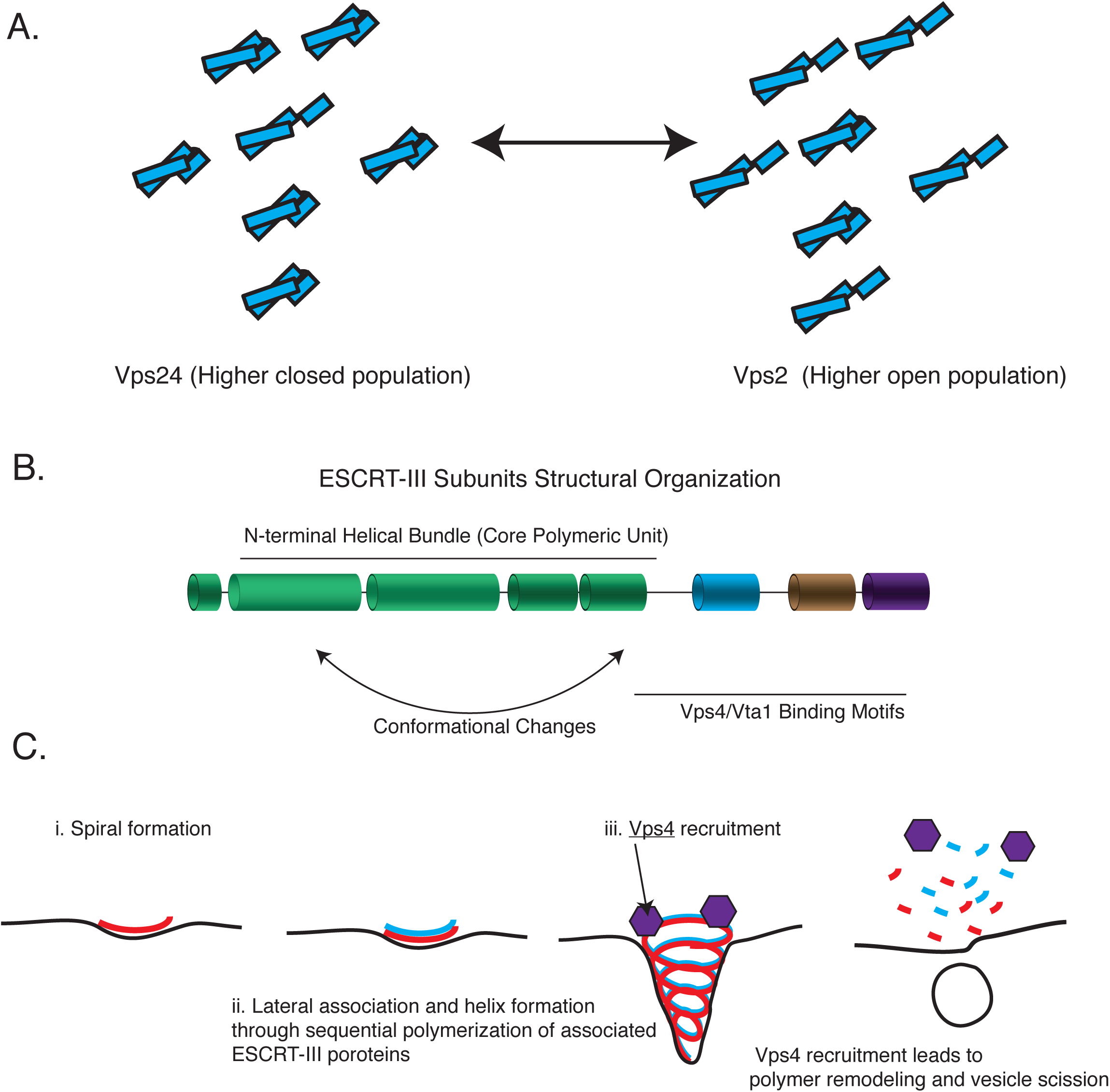
ESCRT-III assembly principles. A) Vps24 and Vps2 are structurally similar proteins that may consist of two different populations of open and closed conformations. Switching between the two conformations mimic each other. B) The domain organization of ESCRT-III subunits and the variousfunctional parts of the structures/sequence. C) The minimal features of ESCRT-III assembly may involve spiral formation, lateral association between copolymers that induce helicity, and recruitment of a disassembly factor such as Vps4.

Following these observations, we propose the following three critical aspects of an ESCRT-III minimal module: (1) a core spiral forming unit (e.g., Snf7), (2) a lateral bundling unit (e.g., Vps24 and Vps2), and (3) an ability to recruit a disassembly machinery (e.g., the AAA+ ATPase Vps4) (Figure 7C).

In addition to the rescue of function phenotypes with Vps24/Vps2, we also find that *did2*Δ can be rescued partially with the overexpression of Vps2. Similarly, swapping the C-termini of Snf7 with that of Vps60 can replace the function of *vps60*Δ. We previously showed that point mutations in Snf7 can rescue the absence of Vps20 (*28*). These data imply, as predicted by evolutionary analyses (*35, 37*), that Vps20-Snf7-Vps60 are more similar to one another, and Vps2-Vps24-Did2 are more alike one another, given the rescue of *vps20*Δ or *vps60* with Snf7 alleles, and *vps24*Δ and *did2*Δ with Vps2 alleles.

*In vitro* reconstitutions suggest that Snf7, Vps24, and Vps2 are essential for membrane budding, and Vps4 for vesicle scission (*11, 17*). Vps4-mediated turnover of a laterally-associating and helix-inducing polymer of Vps24-Vps2 would constrict the Snf7 scaffold to a fission-competent structure, as predicted by simulations (*38*) and as recently proposed to occur in archaeal cell division (*39*). While Did2 and Ist1 are not essential for intraluminal vesicle formation *in vitro* and *in vivo*, they have regulatory roles that are controlled by sequential polymerization dynamics through Vps4 (*17*). So far Vps60 has not been included in *in-vitro* analyses and there have been fewer *in-vivo* analyses on this protein. We find that for Vps60 recruitment to endosomal/vacuole membranes, Vps2 (and likely Vps24) are required. Therefore, Vps60 may be recruited in later stages of polymer formation, in a sequential fashion to the core scaffold, as Did2 does (*17*). Vps60 has been shown to interact with Vta1(*40–42*), and Vta1 is known to be an activator of Vps4 (*40, 43, 44*). Therefore, it is possible that Vps60 is involved in further activating Vps4 function in the later stages of polymer dynamics.

ESCRT-III proteins are integral to all ESCRT-related functions in cells. However, the mechanisms of the specific roles of each ESCRT-III proteins have remained unclear. Based on our work on Vps24 and Vps2 described here and our previous studies on Snf7 and Vps20 (*28*), the molecular features of ESCRT-III subunits should enable future work on rational design of minimal ESCRT-III subunit(s) possessing all properties necessary for intraluminal vesicle formation. Further *in vivo* analyses and *in vitro* reconstitution are required to test whether this minimal ESCRT-III subunit(s) can be created that include the aforementioned features. Some archaeal species consist of only two ESCRT-III proteins, which must possess the minimal properties of ESCRT-III necessary for function (*36*). With the principles learned from our work and from recent studies on ESCRT-III, it will be interesting to study what biochemical features the ancient archaeal ESCRT-III subunits consist, and what additional features were acquired as eukaryotic organelles evolved.

Our earlier understanding of Vps24 and Vps2 suggested that they bound cooperatively to Snf7, but that these were independent proteins with independent and specific functions for MVB sorting. Our data in this study suggest that minor modifications to either one can replace the function of another. These data provide an explanation for why in certain biological processes CHMP3 (mammalian Vps24) may play a minor role (such as in HIV budding), as the isoforms of CHMP2 (Vps2) may already possess the ability to form lateral interactions, and also an ability to recruit the AAA+ ATPase Vps4. The relative contribution of the different ESCRT-III proteins for other ESCRT-dependent processes have not been quantified to the same extent. Further analysis of site-specific ESCRT-III function could allow us to achieve targeted cellular manipulation of ESCRT-dependent processes, understanding of the evolution of these membrane-remodeling polymers and how they contribute to organelle biogenesis.

## Acknowledgments

We thank David Teis for the gift of anti-Vps2 antibody. We thank all members of the Emr lab for discussions. Work in the Emr lab is supported by a Cornell University Research Grant CU3704. Sudeep Banjade is an HHMI fellow of the Damon Runyon Cancer Research Foundation (DRG-2273-16). We are also grateful to the Damon Runyon Cancer Research Foundation for an extension of the fellowship to support our work during the COVID-19 pandemic delays. Shaogeng Tang is a Merck fellow of the Damon Runyon Cancer Research Foundation (DRG-2301-17) on a different project.

## Materials and Methods

### Strains, Plasmids and Reagents

Strains, plasmids and reagents are described in Table 1. Strains previously used were from (*3, 18, 21–23*), and also referenced in the table.

### Random mutagenesis selection

Error prone PCR was used to generate random mutations in the plasmid harboring the S. cerevisiae Vps2 gene. The primers used for this PCR bind the 5’ UTR and the 3’ UTR regions of Vps2. The PCR fragment was transformed into the *vps24*Δ strain in the presence of a linearized Vps2 plasmid by digesting with HindIII and NarI enzymes. The transformants were plated onto plates containing 0.5 μg/mL canavanine, and then replica plated into 4 μg/mL plates. Plasmids were rescued from these colonies that grew on canavanine and then re-transformed into the *vps24*Δ strain to confirm the suppression of *vps24*Δ. Confirmation of *vps24*Δ suppression was done by testing Mup1-pHluorin sorting ability (see below) of the mutants.

### Canavanine Spot Plates

Mid-log cells were serially were diluted to an OD_600_ of 0.1. They were then diluted 10-fold serially, and spot plated in plates containing various concentrations of canavanine. Images of the plates were taken at 3 and 5 days.

### Mup1-pHluorin Flow Cytometry and Immunoblots

Strains harbring Mup1-pHluorin were used to assay endocytosis of this cargo upon methionine addition. Assays were performed as described before (*28*). Briefly, mid-log cells in the presence of synthetic drop-out media were treated with 20 μg/mL L-methionine for 90 minutes and assayed for quenching of pHluorin. Over time as Mup1-pHluorin traffics to the vacuole, fluorescence decreases due to quenching of the pH sensitive pHluorin. Experiments were performed at room temperature and analyses were done on a C6 Accuri flow cytometer from BD Biosciences.

Immunoblots after methionine treatment were performed to analyze free pHlourin, upon degradation of Mup1, as described (*23*). Blots were performed using primary antibody against GFP from Torrey Pines. Imaging of the western blots was performed using an Odyssey CLx imaging system and analyzed using the Image Studio Lite 4.0.21 software (LI-COR Biosciences).

### Doxycycline-mediated Shutdown of the Tet-off Operator

Plasmids (pCM189) used in this study that under the tet-off operator have a CMV promoter and can be regulated by doxycycline titration. For titration experiments, cells were diluted to an OD600 of 0.01. Doxycyline was added at a concentration of 0.25 μg/mL and serially diluted 2-fold over eight times. Cultures were grown until an OD600 of 0.5, and then treated with methionine for Mup1-pHlourin sorting assays or used for co-immunoprecipitation.

### Protein Purification

Vps24, Snf7^R52E^ and Vps2 constructs used in this study were purified as described before (*23*). A combination of affinity (Cobalt Talon resin) and size-exclusion chromatography (SD200increase, GE) were used to purify the proteins. The final buffer under which the proteins are stored was 25 mM Hepes pH 7.5, 150 mM NaCl and 2 mM β-ME. His6-tagged Vps60 and Did2 were purified through cobalt and anion exchange chromatography.

### Electron Microscopy

Lipid monolayers were prepared with a mixture of 60% POPC, 30% POPS and 10% PI3P in chloroform. Carbon-coated electron microscope grids were used to make monolayers and incubate with proteins, as described before(*27*). Grids were stained with 2% ammonium molybdate and imaged on an FEI Morgagni 268 TEM.

### Glycerol Gradient

For Vps24 and Vps24 E114K glycerol gradients, *vps24*Δ was transformed with pCM189 Vps24 or pCM189 Vps24 E114K. 30 ODs of cells expressing these constructs were harvested in PBS buffer. Lysis was performed with PBS buffer, 10% glycerol, 1 mM DTT, Roche protease cocktail and 0.5% Tween-20. Gradient Master 108 from Biocomp was used to make glycerol gradients of 10 to 40%. Centrifugation was performed at 100,000 xg for 4 hours at 4°C. 1 mL fractions were collected from the solutions, TCA precipitated and immunoblotted.

### Fluorescence microscopy

1 mL of mid-log cells was harvested and resuspended in 25 μL water. Imaging was performed on a Deltavision Elite system with an Olympus IX-71 inverted microscope, using a 100X/1.4 NA oil objective. Image extraction and analysis were performed using the FiJi software.

### Sequence and structural analyses

Mafft(*45*) and Jalview (*46*) were used to analyze sequences. Heliquest was used for helical wheel analysis(*47*). Structural models were made using UCSF Chimera(*48*).

### Co-immuoprecipitation

30 ODs of mid-log cells were harvested and washed with cold MilliQ H2O, and resuspended in 1 mL phosphate-saline buffer (PBS), 10% glycerol, 1 mM DTT and 1 mM EDTA, including protease inhibitor cocktail from Roche. Lysis was performed by bead-beating (Zirconia-Silicon beads) twice for 30 s, with 30s intervals on ice. Lysate was treated with 1% Triton X-100 and rotated for 20 minutes. Lysate was cleared by centrifugation at 16,000 xg at 4°C. The supernatant was treated with protein G beads (Dynabeads) for 30 min at 4°C to remove nonspecific binding. The magnetic beads used for this assay were allowed to settle with a magnetic Eppendorf-tube rack, and the supernatant was applied with 1/250 v/v of anti-Snf7 antibody. After 1-hour incubation at 4°C, the beads were washed twice with 20X fold bead volume of the lysis buffer. Proteins were eluted by incubating the beads at 65°C for 10 minutes in sample buffer (150 mM Tris-Cl, pH 6.8, 8 M urea, 10% SDS, 24% glycerol, 10% v/v □ME, and bromophenol blue). Anti-Snf7 and anti-Vps2 antibodies were used to probe for eluted proteins through Western blots.

---

**Figure 1 - Fig. Supp. 1.**
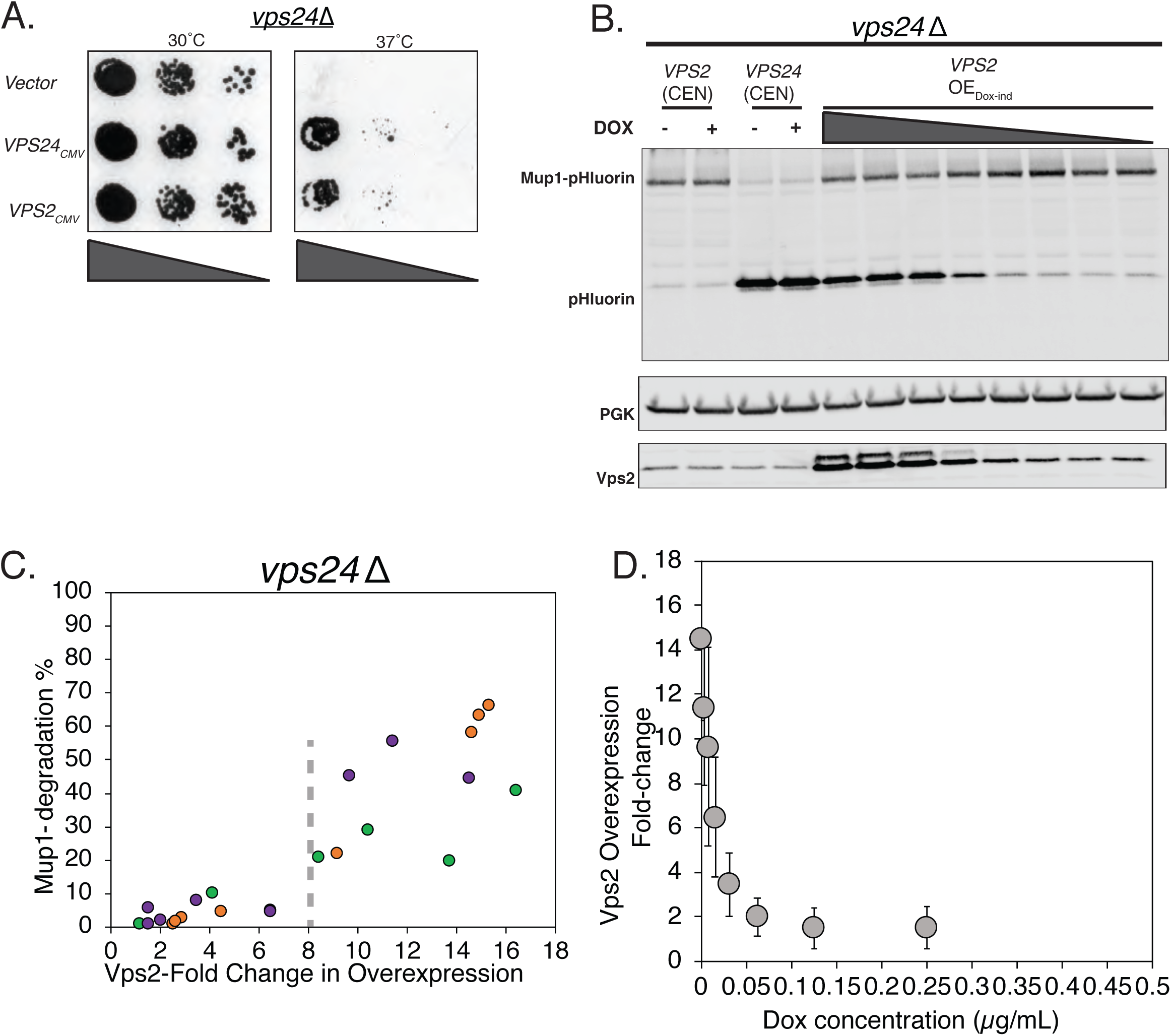
Vps2 overexpression can rescue the defect of *vps24*Δ. A)Vps2 overexpression with a CMV-promoter/Tet-operator rescues the temperature sensitivity defect of *vps24*Δ. B) Immunoblot of pHLuorin showing the cleavage of Mup1-pHluorin after 90 minutes of methionine addition. Vps2 was expressed either in a single-copy centromeric (CEN) plasmid or under a doxycycline-inducible CMV promoter. Expression of Vps2 was controlled by titrating the concentration of doxycycline. C) Mup1-sorting characterization with changes in Vps2 expression level. The different colors represent different set titration experiments. D) Plot showing the control of expression levels with doxycycline titration.

**Figure 1 - Fig. Supp. 2.**
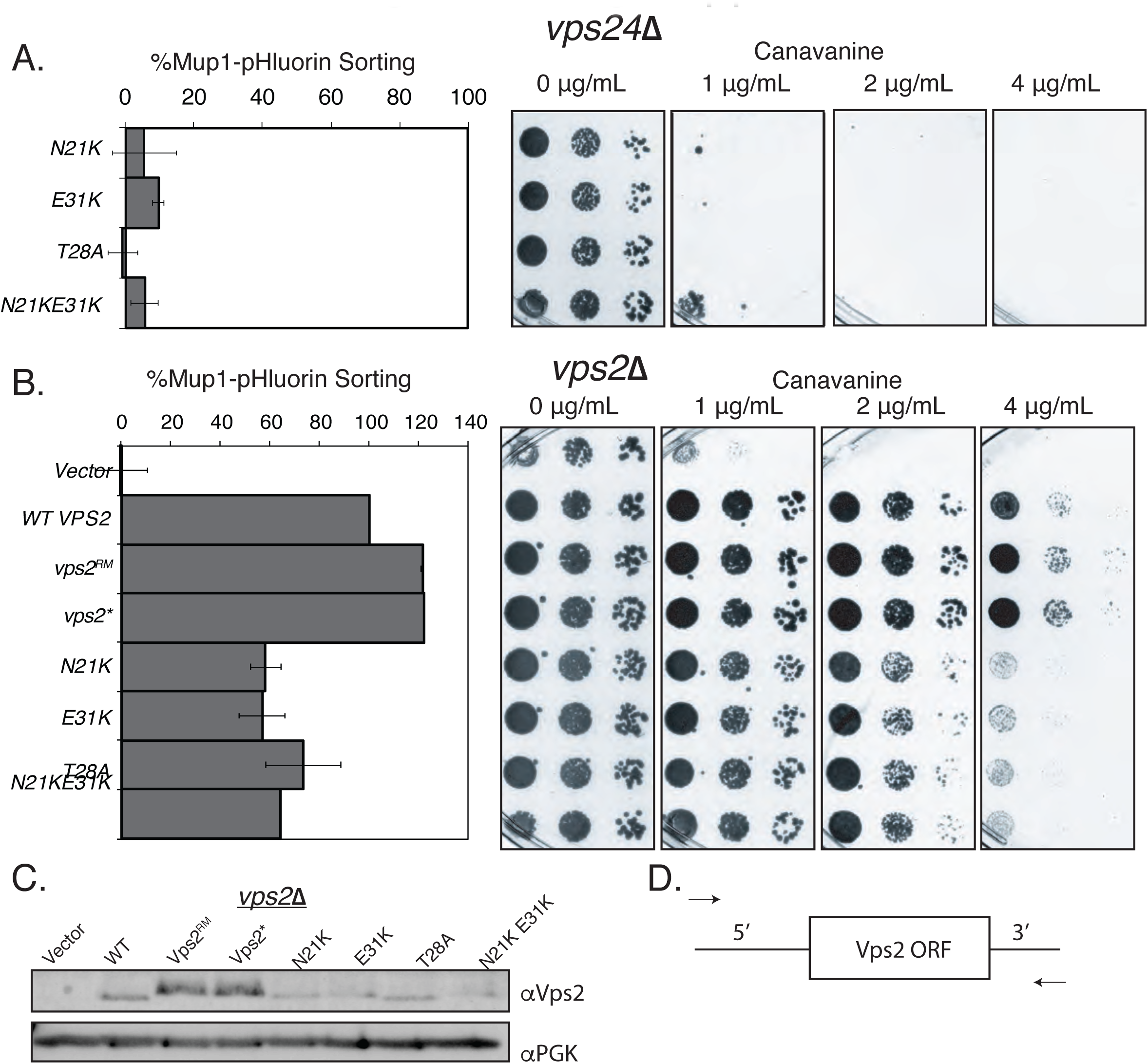
Vps2 N-terminal mutations can rescue the defect of *vps24*Δ. A) Flow-cytometry for Mupl-pHluorin sorting and canvanine sensitivity assay in *vps24*Δ for the N-terminal helix-1 mutations in Vps2 (compare with Fig. 1 C). In these constructs the promoters were endogenous, WT promoters. B) In a *vps2*Δ background, the suppressors *vps2^RM^* and *vps2\** also have higher sorting capabilities in both Mupl-sorting assay and in canavanine sensitivity assay. The promoter regions of *vps2^RM^* and *vps2\** contains mutations, but other constructs are with WT promoters. C) Immunoblots of various Vps2 mutants, the same constructs as used in Fig. 1C, and Fig. 1 - Supp. 2 A and C. D) Design of the randomly mutagenized Vps2 plasmid - primers bind to the 5’ and 3’ UTR regions of Vps2.

**Figure 1 - Fig. Supp. 3.**
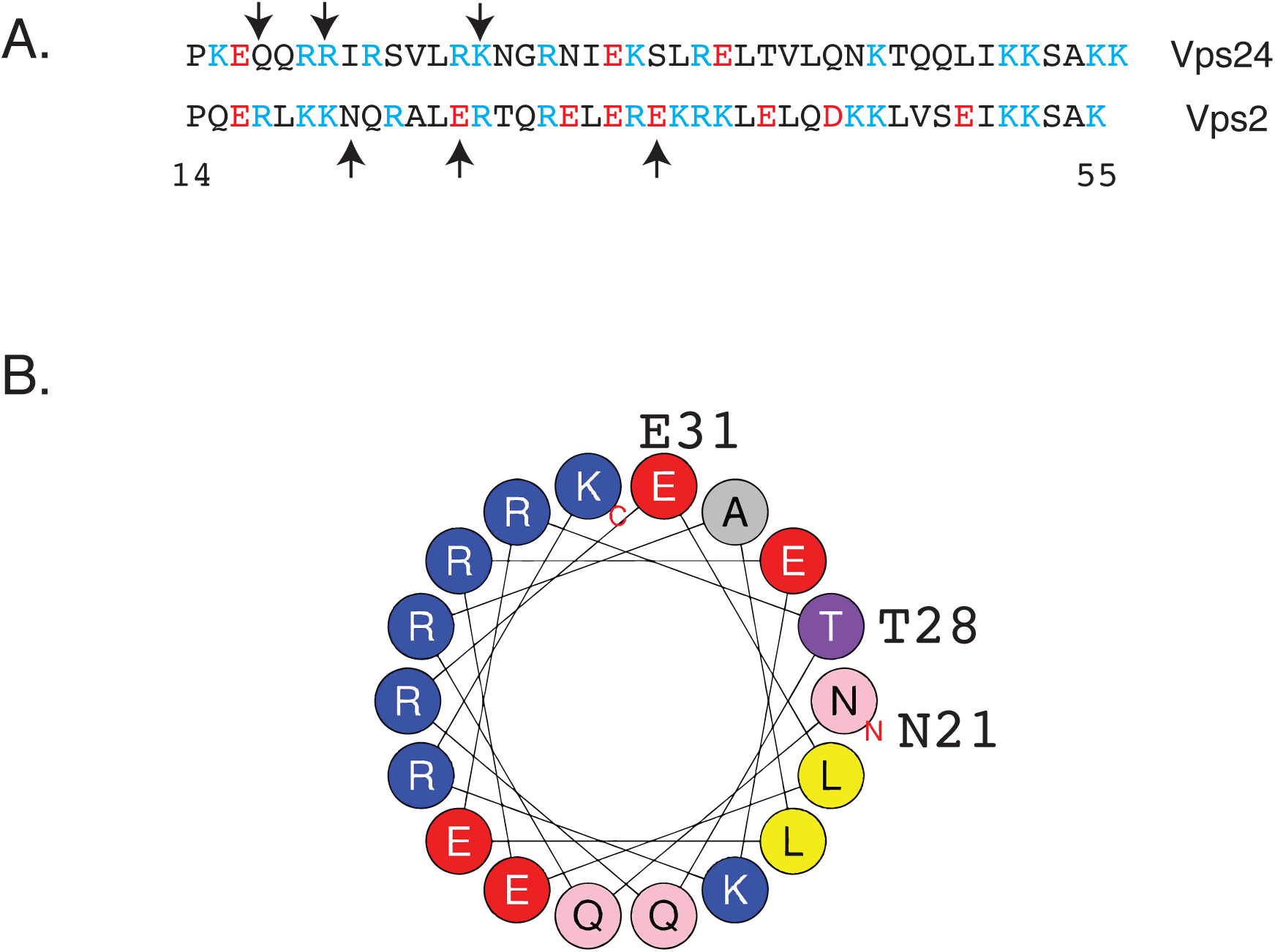
Helix-1 region of Vps2 is important for binding to Snf7. A) Sequence alignment of helices 1 of Vps24 and Vps2. Cyan-colored residues are basic amino acids, and red colors represent acidic amino acids. Arrows in Vps24 sequence point to mutations that rescue the defect of the *snf7D^131K^* allele (Banjade et al., 2019). Arrows in Vps2 sequence represent the location of the mutations that rescue *vps24*Δ. B) Helical wheel representation of part of the helix-1 region of Vps2 (Heliquest).

**Figure 3 - Fig Supp. 1.**
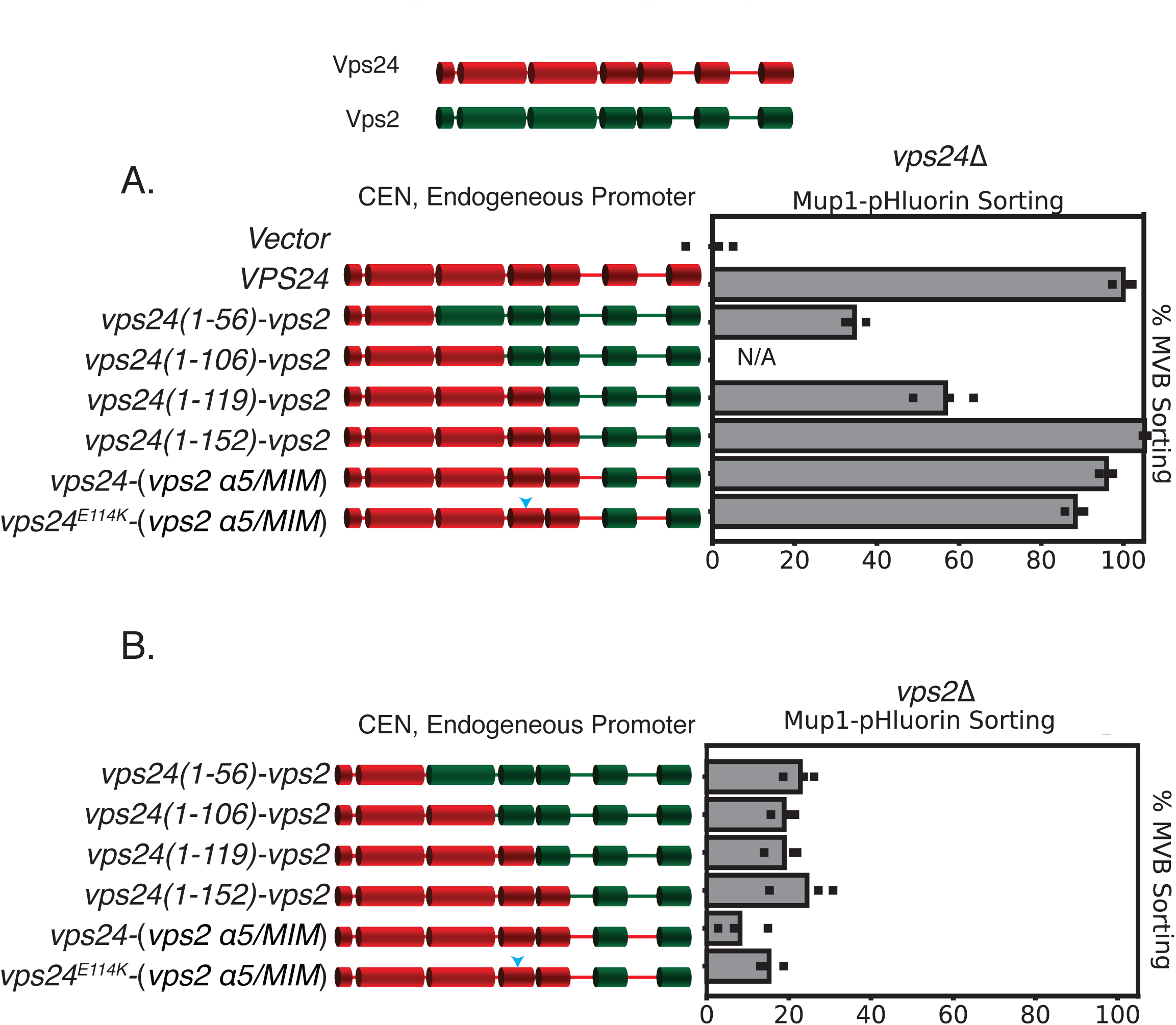
Chimeras of Vps24-Vps2 are functional proteins. Top figure depicts the domain organization of Vps2 and Vps24. A) Mup-pHluoring sorting assay with several chimeras of Vps24-Vps2, showing that the replacement of the C-terminal regions of Vps2 onto Vps24 keeps the constructs functional. B) The same constructs as in (A) do not supress *vps2*Δ, as they are under endogeneous prometers. “CEN” represents denotation for centro-meric plasmid.

**Figure 3 - Fig Supp. 2.**
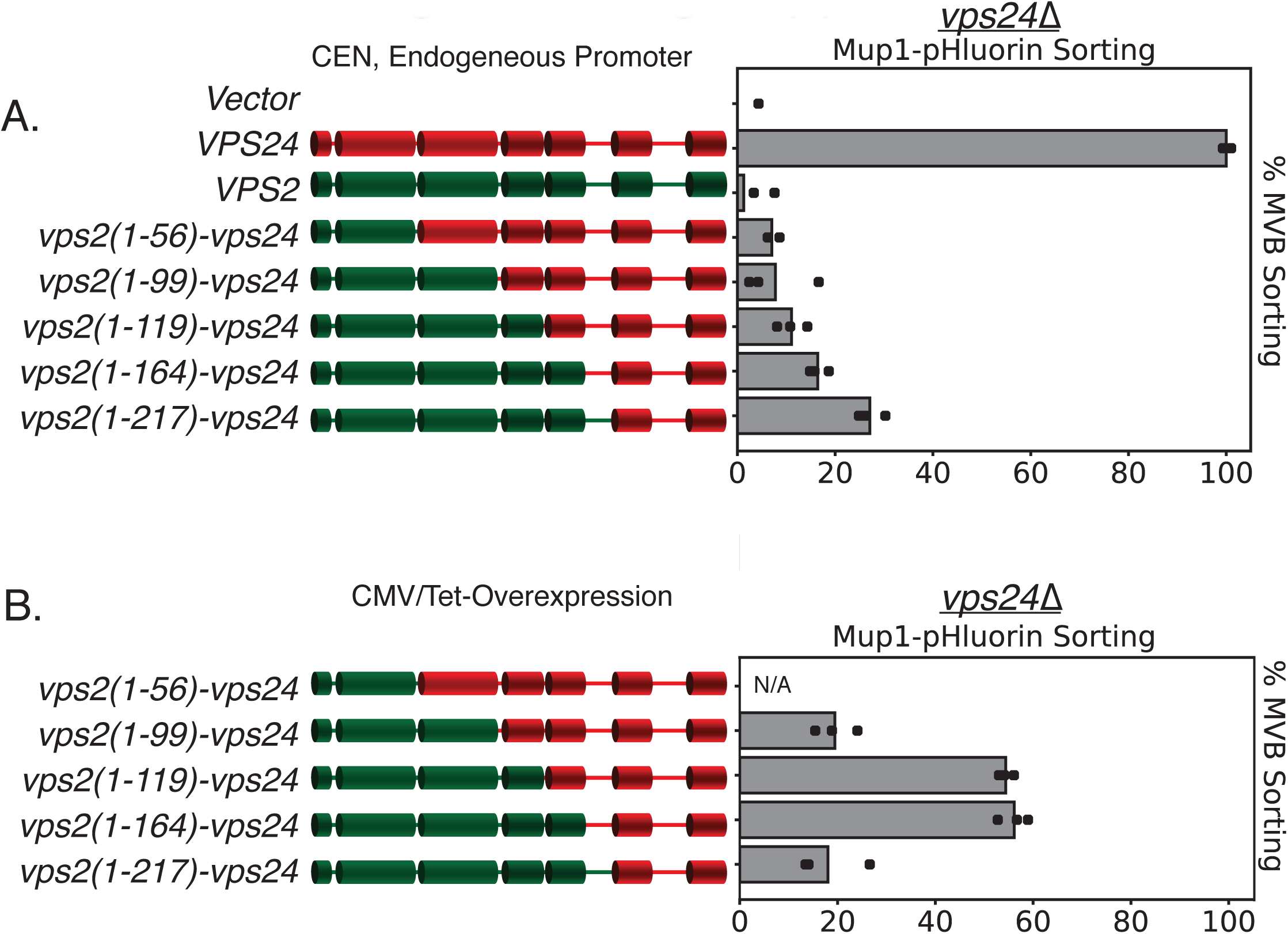
Simple modifications in Vps24 can be made to mimic Vps2. A) Under the endogenous promoter and centromeric plasmid (CEN), various chimeras of Vps24-Vps2 do not support the sorting of Mup1-pHluorin, but some of the same constructs when overexpressed can rescue *vps24*Δ (B). Note that the N-terminus of Vps2 needs to be intact to mimic Vps24. Overexpression (~16 fold) was achieved with a CMV-promoter, Tet-operator system.

**Figure 4- Fig. Supp 1.**
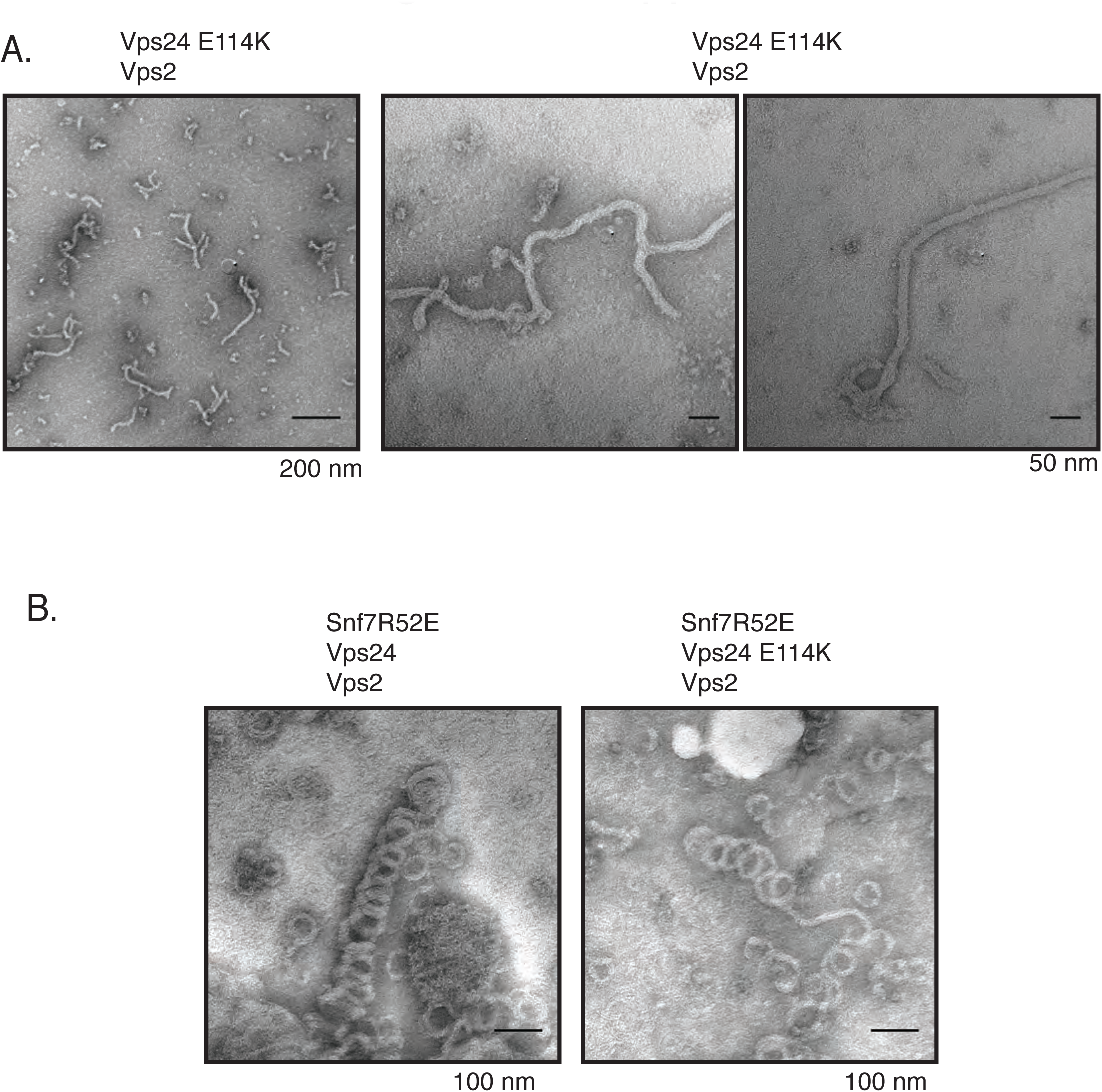
Vps24 (E114K) associates with Vps2. A) Electron microscopy images of Vps24E114K assembled with Vps2 at concentrations of 5*μ*M each. Two images on the right are zoomed-in images of the same polymers. Vps24 E114K alone or Vps24 with Vps2) do not form such polymers. B) Vps24E114K mutant with Vps2 and Snf7-R52E still form 3D helices. Snf7-R52E is a mutant that has a lower critical concentraiton for polymerization as a higher fraction of this protein is in an open conformation (Henne et al., 2012).

**Figure 4- Fig. Supp 2.**
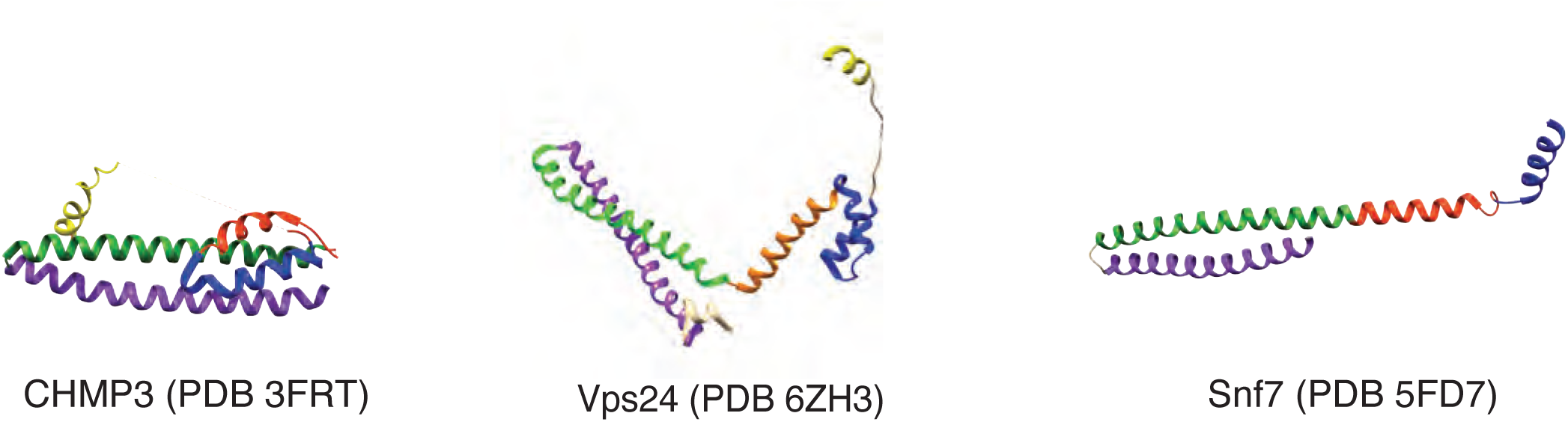
Structures of the autoinhibited CHMP3, and the filament forming conformations of Vps24 and Snf7.

**Figure 5 - Figure Supplement 1.**
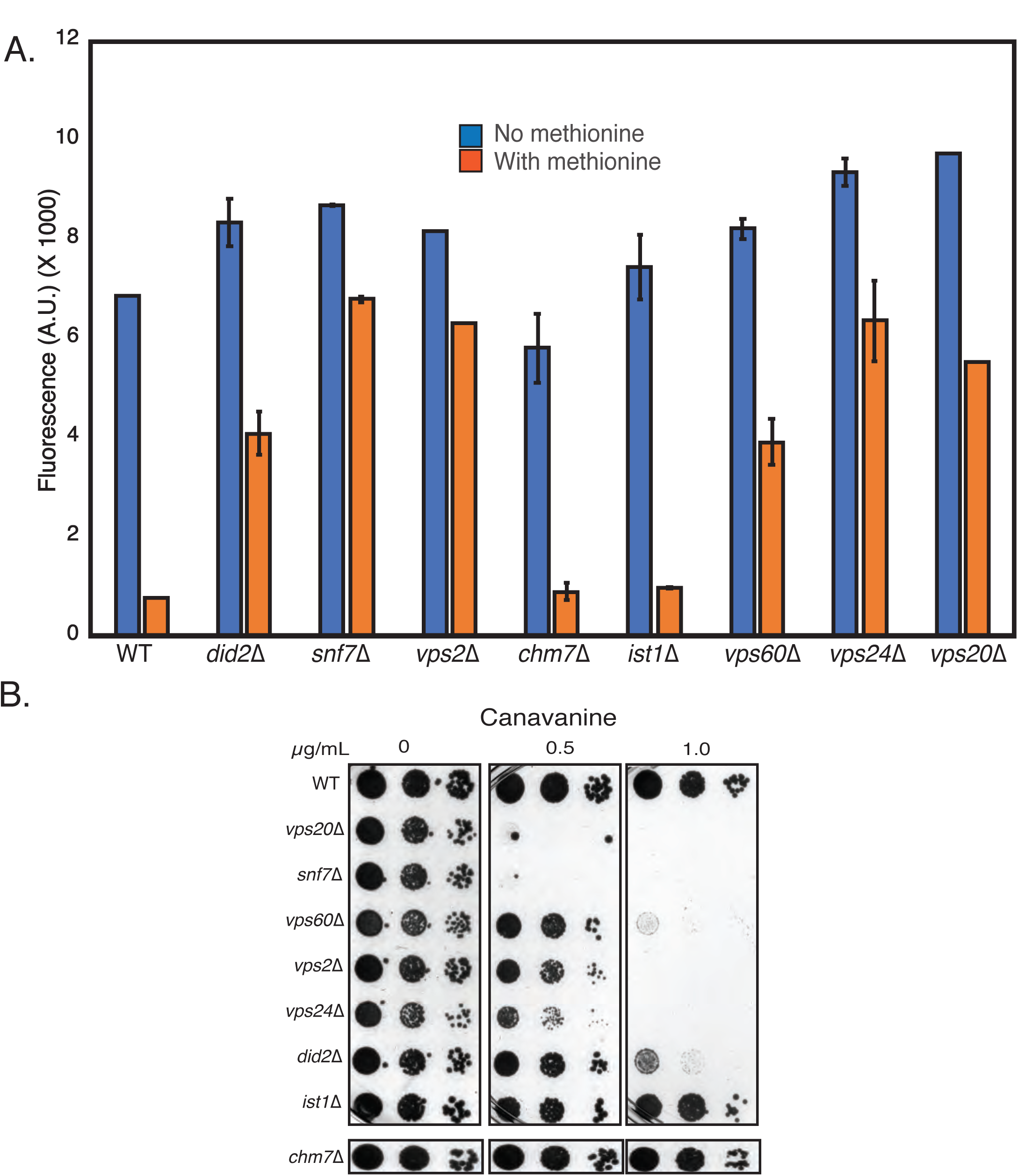
A) Relative effects of all ESCRT-III mutants for defects in cargo sorting, using Mup1-pHluorin assay. Fluorescence of 100,000 cells were measured after 90 minutes of adding of 20 *μ*g methionine. B) Canavanine sensitivity assays of the ESCRT-III mutants.

**Figure 6 - Fig. Supp. 1.**
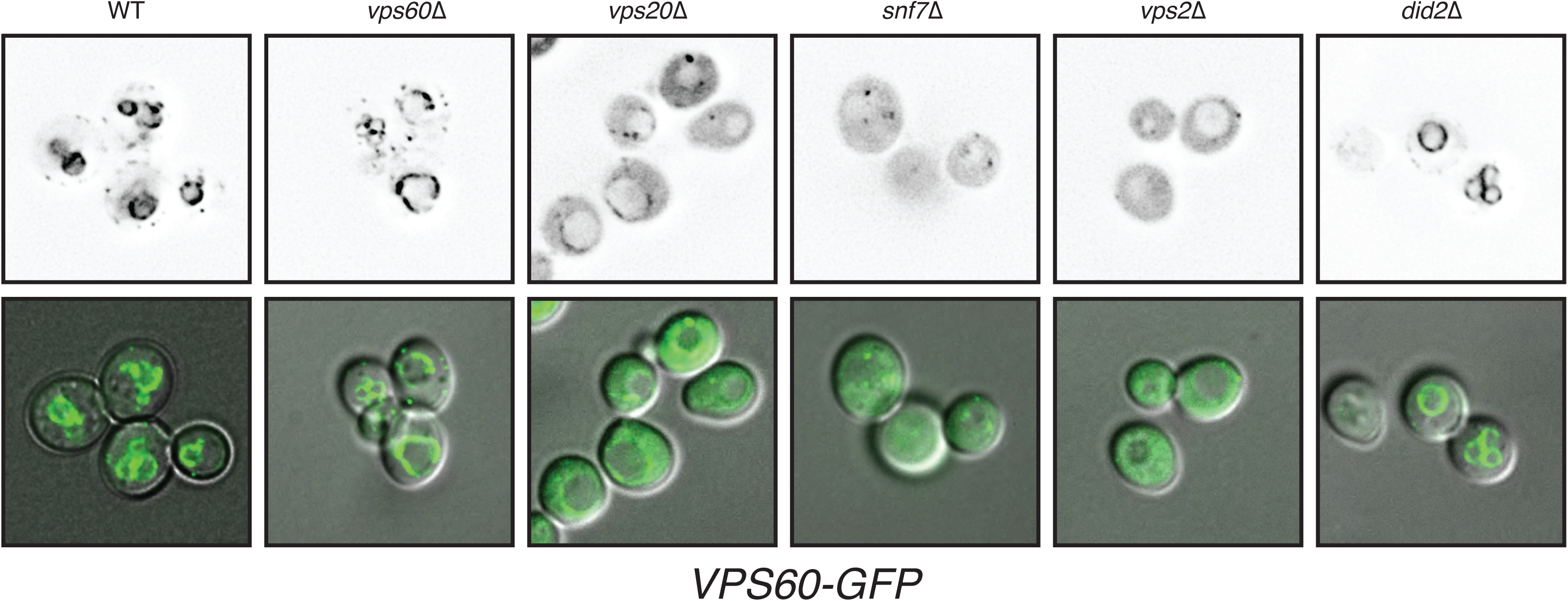
Localization of Vps6O-GFP in different ESCRT-III mutants. Top panel represents gray-scaled GFP fluorescence and bottom panel represents a merge of GFP and DIC channels. While in WT strains *VPS60-GFP* primarily localizes to punctae (membranes), in various mutants the cytoplasmic signal is increased.

